# The heat shock transcription factor HSF-1 protects *Caenorhabditis elegans* from peroxide stress

**DOI:** 10.1101/2020.07.13.200634

**Authors:** Francesco A. Servello, Javier Apfeld

## Abstract

Cells induce conserved defense mechanisms that protect them from oxidative stress. How these defenses are regulated in multicellular organisms is incompletely understood. Using the nematode *Caenorhabditis elegans*, we show that the heat shock transcription factor HSF-1 protects the nematode from the oxidative stress induced by environmental peroxide. In response to a heat shock or a mild temperature increase, HSF-1 protects the nematodes from subsequent oxidative stress in a manner that depends on HSF-1’s transactivation domain. At constant temperature, HSF-1 protects the nematodes from oxidative stress independently of its transactivation domain, likely by inducing the expression of *asp-4/cathepsin D* and *dapk-1/dapk*. Thus, two distinct HSF-1-dependent processes protect *C. elegans* from oxidative stress.

## Introduction

Organisms have evolved conserved defense mechanisms that enable survival in adverse environments including a wide variety of physical, chemical, and biological stressors [1], such as heat [2,3], radiation [4], oxygen levels [5], and feeding [6]. Induction of these defenses can lead to lasting protection in both harmful and normal environments, a phenomenon called hormesis [7]. In the nematode *C. elegans*, one of the ways that hormesis can be triggered is by a brief exposure to high heat (heat shock). Heat shock induces a heat shock response, which protects *C. elegans* from subsequent lethal heat exposure [2,8], from subsequent infection [9], and increases lifespan in the absence of stress [2].

The heat shock response is a conserved transcriptional response, mediated in part by the heat shock transcription factor HSF-1/HSF, that protects animals from heat stress [10]. In *C. elegans*, HSF-1 activates the expression of protective chaperones and other heat shock proteins [11–13] via a mechanism that requires the transactivation domain of HSF-1 [11,14–16]. More recently, HSF-1 was found to activate the expression of a different set of heat-stress protective genes by a mechanism that does not require its transactivation domain [17]. Here, we show that HSF-1’s transactivation domain is required for heat stress to protect *C. elegans* from subsequent oxidative stress. In addition, we show that in the absence of heat stress, HSF-1 protects *C. elegans* from oxidative stress independently of its transactivation domain. This effect is likely mediated by two transactivation-domain independent targets of HSF-1, ASP-4/Cathepsin and DAPK-1/DAPK. Our work advances the understanding of how heat stress and HSF-1 influence resilience to oxidative stress in *C. elegans*.

## Results

We speculated that the adaptive response triggered by heat shock may also protect against oxidative stress. To test that hypothesis, we determined the extent to which heat shock influenced the nematode’s resistance to environmental peroxide. We exposed late L4 stage nematodes cultured at 20°C to 34°C for one or two hours, let the animals recover at 20°C for two days, and subsequently measured their survival to 6 mM tert-butyl hydroperoxide (tBuOOH) at 20°C. Wild type nematodes heat-shocked for one hour survived 17% longer than control nematodes that were not heat-shocked (Figure 1A). Nematodes heat-shocked for two hours survived 25% longer than those heat shocked for one hour (Figure 1A). We conclude that heat shock leads to a subsequent dose-dependent increase in the nematode’s peroxide resistance.

**Figure 1.**
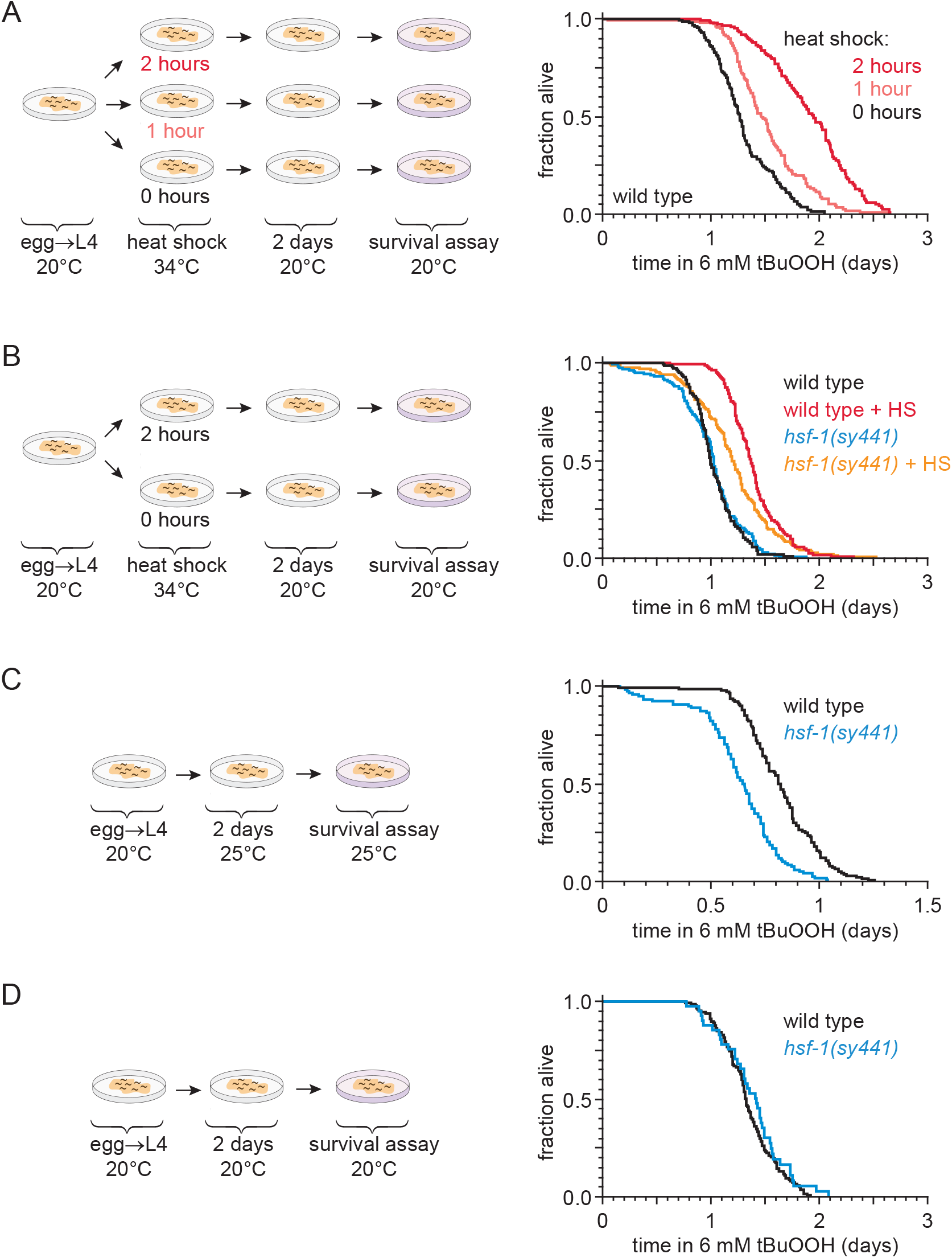
HSF-1 regulates oxidative stress resistance in response to temperature changes dependent on its transactivation domain. (A) Prior heat shock (HS) increased peroxide resistance in a dose-dependent manner in wild-type nematodes cultured at 20°C. Nematodes were heat shocked at the L4 stage and assayed for survival in the presence of 6 mM tert-butyl hydroperoxide (tBuOOH) on day 2 of adulthood. (B) The increase in peroxide resistance induced by HS was partially suppressed by the *hsf-1(sy441)* mutation, which removes the HSF-1 transactivation domain. (C) *hsf-1(sy441)* decreased peroxide resistance when nematodes were shifted from 20°C to 25°C at the L4 larval stage. (D) *hsf-1(sy441)* did not affect peroxide resistance when nematodes were cultured continuously at 20°C. The diagrams summarize the experimental strategy leading to each assay. Statistical analyses are in Supplementary Table 1.

Heat shock induces resistance to subsequent lethal heat exposure partially through the action of the heat shock transcription factor HSF-1 [18]. We set out to determine whether HSF-1 is also required for heat shock to induce subsequent peroxide resistance. The *hsf-1(sy441)* mutation is a premature stop codon that truncates the HSF-1 protein, leaving the DNA binding and multimerization domains intact, but removing the transactivation domain required for HSF-1 to induce the expression of chaperones and other heat shock proteins upon heat shock [11,17]. In the absence of heat shock, the peroxide resistance of *hsf-1(sy441)* mutants and wild type nematodes was indistinguishable (Figure 1B). Therefore, the transactivation domain of HSF-1 did not appear to regulate peroxide resistance in nematodes cultured continuously at 20°C. In contrast, heat-shocked *hsf-1(sy441)* mutants exhibited a smaller increase in peroxide resistance than heat-shocked wild type nematodes (Figure 1B). We conclude that heat shock increases subsequent peroxide resistance in part via a process that requires HSF-1-dependent transactivation.

HSF-1-transactivation dependent gene expression can also be induced in response to much milder temperature shifts, including induction in response to an up-shift from 15°C to 25°C [19]. To determine whether HSF-1-transactivation promotes peroxide resistance at higher growth temperatures, we determined the peroxide resistance of wild type and *hsf-1(sy441)* mutants cultured at 20°C until the onset of adulthood, and then shifted to 25°C for two days. When the nematodes were up-shifted to 25°C, the *hsf-1(sy441)* mutation caused a decrease in peroxide resistance relative to wild type animals (Figure 1C), but when they remained at 20°C *hsf-1(sy441)* did not affect peroxide resistance (Figure 1D). We conclude that HSF-1’s transactivation domain is necessary to increase peroxide resistance when nematodes are up-shifted to 25°C, but not when grown at 20°C.

Because HSF-1-transactivation did not influence peroxide resistance of nematodes cultured at 20°C, we were surprised to find that, at 20°C, *hsf-1(sy441)* mutants overexpressing an *hsf-1* C-terminal truncation allele missing the transactivation domain [17] exhibited a three-fold increase in peroxide resistance (Figure 2A). This indicated that HSF-1 could influence peroxide resistance at 20°C in a transactivation-domain independent manner. Nematodes overexpressing full-length *hsf-1(+)* survived three-fold longer than wild type nematodes at 20°C (Figure 2B), indicating that full-length HSF-1 can be sufficient to increase peroxide resistance. Peroxide resistance was also increased by the *cg116* null mutation in *hsb-1* (Figure 2C), a negative regulator of HSF-1 activity [20,21], suggesting that increased HSF-1 activity is sufficient to increase peroxide resistance at 20°C. To determine whether full-length HSF-1 normally increases peroxide resistance, we knocked down *hsf-1* via RNA interference (RNAi) and determined the nematode’s peroxide resistance. RNAi of *hsf-1* decreased peroxide resistance by 50% at 20°C (Figure 2D). We propose that HSF-1 normally functions independently of its transactivation domain to increase peroxide resistance at 20°C.

**Figure 2.**
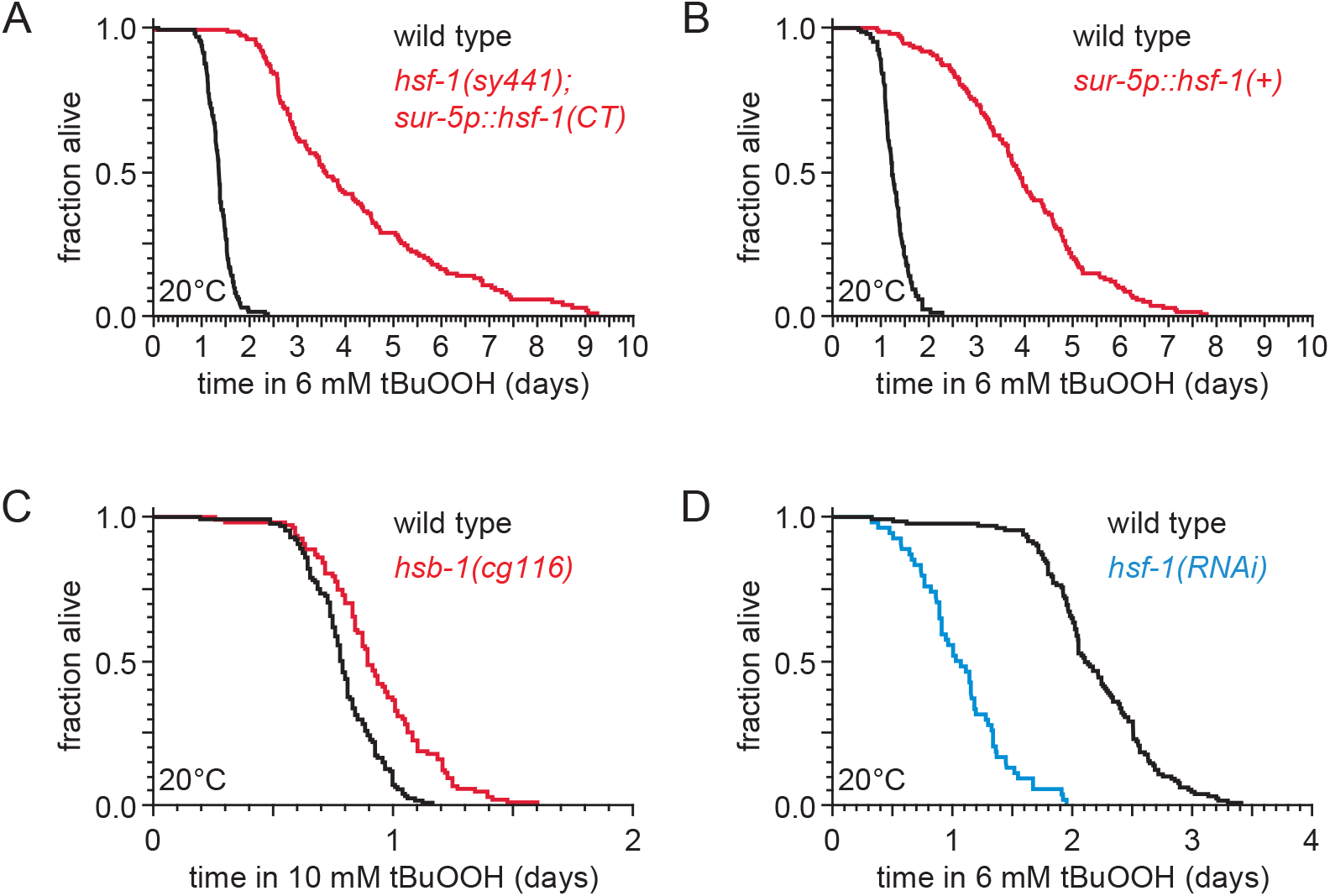
HSF-1 regulates oxidative stress resistance independently of its transactivation domain when nematodes are cultured continuously at 20°C. (A) Peroxide resistance was increased in *hsf-1(sy441)* mutants overexpressing an *hsf-1* (B) Peroxide resistance was increased by overexpression of a full-length *hsf-1(+).* (C) *hsb-1(cg116)* increased peroxide resistance. (D) Knockdown of *hsf-1* via RNAi decreased peroxide resistance. Statistical analyses are in Supplementary Table 2.

Finally, we explored how HSF-1 may increase peroxide resistance in a transactivation-domain independent manner. Transcriptomic and proteomic analyses identified 98 genes whose expression can be increased by HSF-1 in either a transactivation-dependent manner or a transactivation-independent manner [17]. We focused on the potential role of two of those genes, *asp-4* and *dapk-1*, because knockdown of each of these genes did not cause developmental arrest nor affect survival to lethal heat stress [17]. *asp-4* encodes an aspartyl protease related to cathepsins D and E required for neurodegeneration [22]. *dapk-1* encodes a death-associated serine/threonine protein kinase involved in innate immune responses to wounding [23]. *asp-4(ok2693)* deletion mutation and the *dapk-1(ju4)* loss-of-function mutation both lowered peroxide resistance of nematodes cultured at 20°C (Figure 3A-B). We propose that HSF-1 increases peroxide resistance in a transactivation domain independent manner by increasing the expression of ASP-4/Cathepsin and DAPK-1/DAPK.

**Figure 3.**
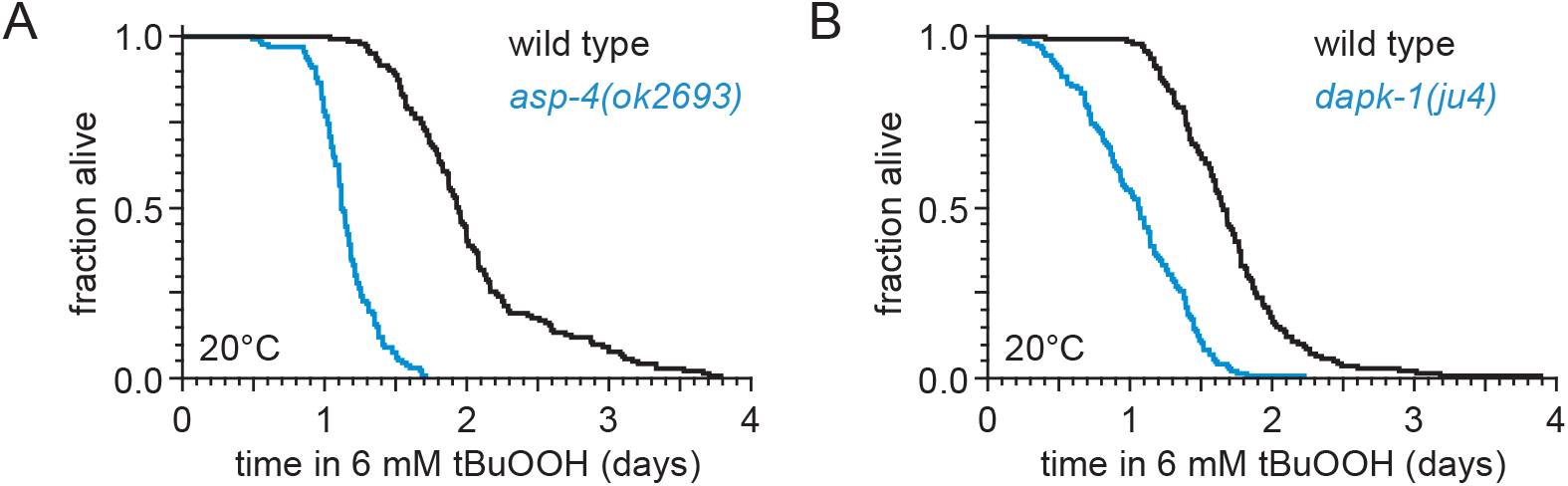
Genes induced by HSF-1 independently of its transactivation domain regulate oxidative stress resistance when nematodes are cultured continuously at 20°C. (A) *asp-4(ok2693)* decreased peroxide resistance. (B) *dapk-1(ju4)* decreased peroxide resistance. Statistical analyses are in Supplementary Table 3.

## Discussion

We propose that HSF-1 induces two partially overlapping sets of genes to protect *C. elegans* against oxidative stress: one set is induced in response to heat shock or mild temperature upshift in a manner that requires HSF-1’s transactivation domain and the other set is induced at a constant favorable temperature in a manner that does not require that domain (Figure 4). The induction of these gene sets by the two types of HSF-1-dependent processes may enable *C. elegans* to tune its defenses to varying environments. Peroxide resistance is increased by *asp-4/Cathepsin* and *dapk-1/DAPK*, two genes that HSF-1 can induce with or without its transactivation domain; possibly to activate protein degradation [22] or innate immune responses [23]. Interestingly, knockdown of *asp-4* or *dapk-1* did not affect survival at 34°C [17], suggesting that HSF-1 regulates peroxide and heat resistance by separable mechanisms. It would be interesting to determine the extent to which HSF-1 can regulate both the basal level of peroxide resistance and the increase in peroxide resistance induced by changes in environmental temperature through the action of a common set of targets.

**Figure 4.**
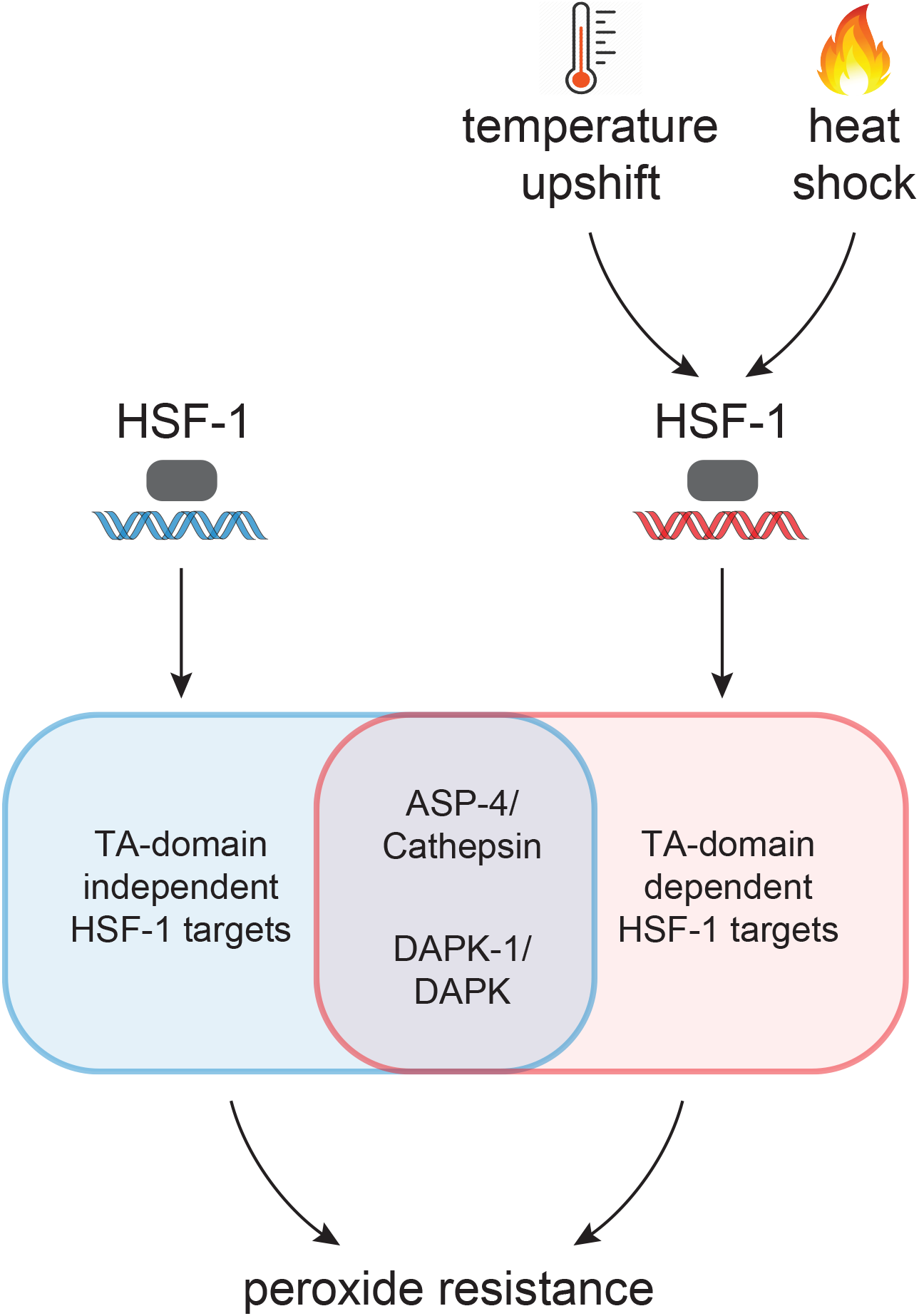
HSF-1 plays multiple roles in the regulation of oxidative stress resistance. Model: Two HSF-1-dependent processes protect against the oxidative stress caused by environmental peroxide in nematodes grown at constant temperature and in nematodes that experienced a temperature upshift or a heat shock. “TA-domain” is the HSF-1 transactivation domain.

## Materials and Methods

### *C. elegans* culture, strains, and transgenes

*C. elegans* were cultured on NGM agar plates seeded with *E. coli* OP50 using standard methods. The following strains were used for this study: QZ0 Bristol N2, QZ112 *hsf-1(sy441) I*, AGD710 *uthIs235[sur-5p::hsf-1::unc-54 3’UTR + myo-2p::tdTomato::unc-54 3’ UTR]*, AGD794 *hsf-1(sy441) I; uthIs225[sur5p::hsf-1(CT-Delta)::unc-54 3’UTR + myo-2p::tdTomato::unc-54 3’ UTR]*, CH116 *hsb-1(cg116) IV*, RB2035 *asp-4(ok2693) X*, and CZ419 *dapk-1(ju4) I*.

### Survival assays

Nematodes were at 20°C, except in temperature up-shift experiments, where they were at 20°C until the onset of adulthood, and then at 25°C. Heat shock treatment was performed in late L4 larvae by exposure to 34°C as described [25]. RNAi treatment was performed by culturing nematodes and their parents on petri plates seeded with *E. coli* HT115 (DE3) expressing double stranded RNA targeting *hsf-1* from the Ahringer library [26] at an OD600 of 20 on NGM plates supplemented with 2mM isopropyl β-d-1-thiogalactopyranoside. Empty vector plasmid pL4440 was used as control. During adulthood, nematodes were cultured and then assayed for survival on plates with 10 μg/ml 5-fluoro-2-deoxyuridine (Sigma), to avoid vulval rupture [27] and eliminate live progeny. Survival to 6mM or 10 mM tert-butyl hydroperoxide (Sigma) was determined on day 2 adults using a *C. elegans* lifespan machine scanner cluster [28] as described [29]. A typical experiment consisted of up to four genotypes or conditions, with 4 assay plates of each genotype or condition, each assay plate containing a maximum of 40 nematodes, and 16 assay plates housed in the same scanner.

### Statistical analysis

Statistical analyses were performed in JMP Pro version 14 (SAS). Survival curves were calculated using the Kaplan-Meier method. We used the log-rank test to determine if the survival functions of two or more groups were equal.

## Acknowledgements

We thank Erin Cram, Charlotte Kelley, Jodie Schiffer, and Yuyan Xu for critical reading and detailed comments on our manuscript. We are grateful to Nicholas Stroustrup for his generous help and for continuing to develop and support the Lifespan Machine software. We benefitted from discussions with members of Javier Apfeld’s and Erin Cram’s lab. Some strains were provided by the CGC, which is funded by NIH Office of Research Infrastructure Programs (P40 OD010440). The research was supported by National Science Foundation CAREER grant 1750065 to J.A. and a Northeastern University Tier 1 award to J.A.

## Competing interests

The authors declare that no competing interests exist.

---

**Supplementary table 1.**
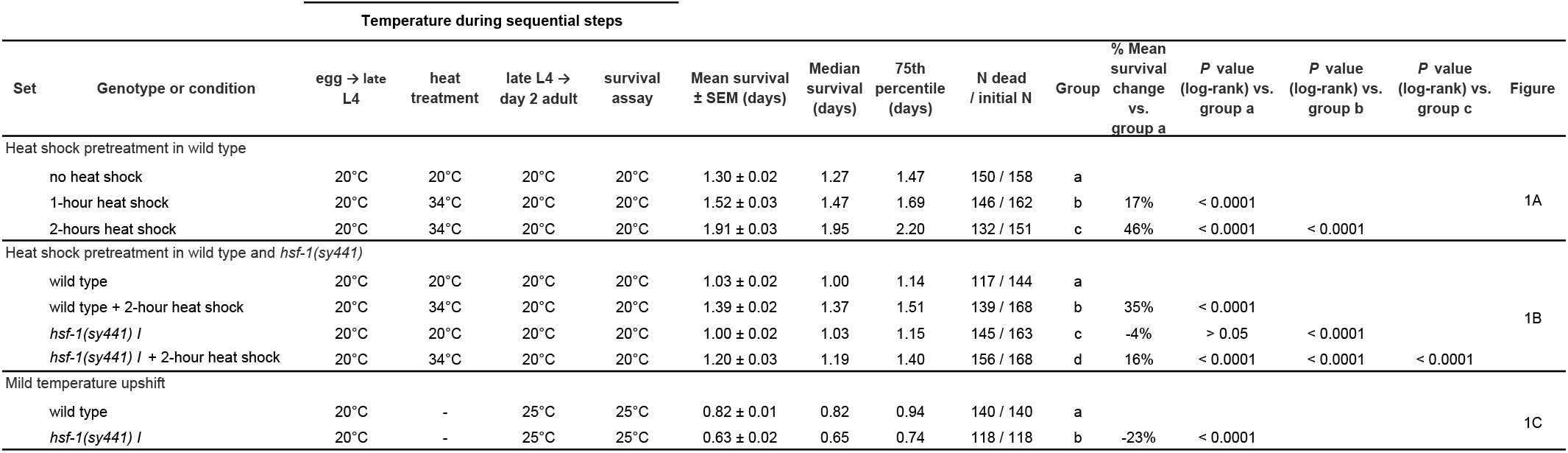
Statistical analysis for Figure 1.

**Supplementary table 2.**
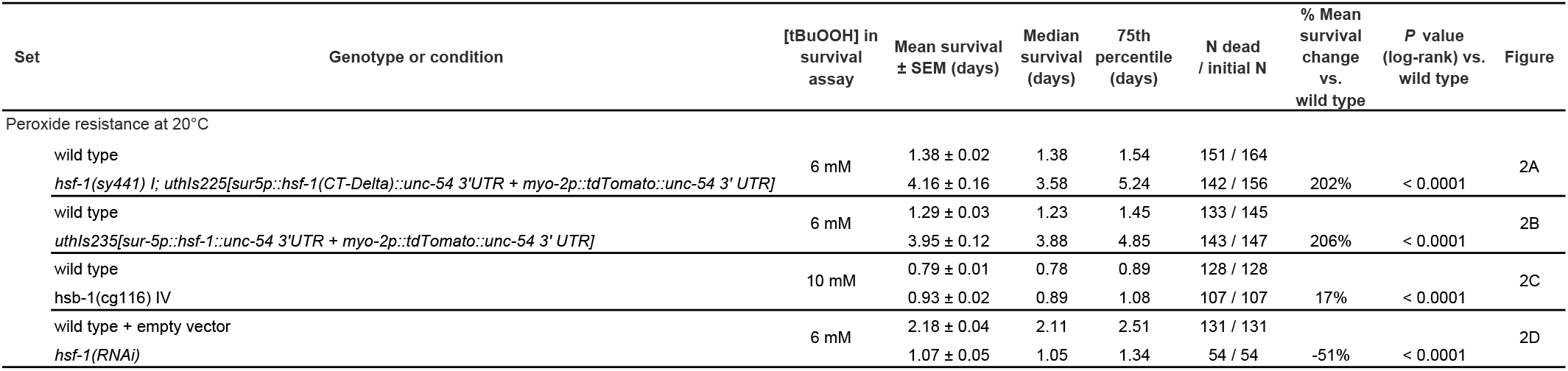
Statistical analysis for Figure 2.

**Supplementary table 3.** Statistical analysis for Figure 3.

| Set | Genotype or condition | Mean survival<br>± SEM (days) | Median<br>survival<br>(days) | 75th<br>percentile<br>(days) | N dead<br>/ initial N | % Mean<br>survival<br>change<br>vs.<br>wild type | P value<br>(log-rank) vs.<br>wild type | Figure |
| --- | --- | --- | --- | --- | --- | --- | --- | --- |
| Peroxide resistance at 20°C |  |  |  |  |  |  |  |  |
|  | wild type | 2.04 ± 0.05 | 1.94 | 2.21 | 142 / 142 |  |  | 3A |
|  | *asp-4(ok2693) X* | 1.15 ± 0.02 | 1.12 | 1.25 | 133 / 133 | -44% | < 0.0001 |  |
|  | wild type | 1.69 ± 0.04 | 1.66 | 1.88 | 140 / 140 |  |  | 3B |
|  | *dapk-1(ju4) I* | 1.04 ± 0.03 | 1.06 | 1.38 | 145 / 145 | -38% | < 0.0001 |  |

## Notes

### Competing Interest Statement

The authors have declared no competing interest.

